# Reconstruction and visualization of large-scale volumetric models of neocortical circuits for physically-plausible in silico optical studies

**DOI:** 10.1101/164483

**Authors:** Marwan Abdellah, Juan Hernando, Nicolas Antille, Stefan Eilemann, Henry Markram, Felix Schürmann

**Author notes:** This article is accepted for publication in BMC Bioinformatics 2017.

## Abstract

**Background** We present a software workflow capable of building large scale, highly detailed and realistic volumetric models of neocortical circuits from the morphological skeletons of their digitally reconstructed neurons. The limitations of the existing approaches for creating those models are explained, and then, a multi-stage pipeline is discussed to overcome those limitations. Starting from the neuronal morphologies, we create smooth piecewise watertight polygonal models that can be efficiently utilized to synthesize continuous and plausible volumetric models of the neurons with solid voxelization. The somata of the neurons are reconstructed on a physically-plausible basis relying on the physics engine in Blender.

**Results** Our pipeline is applied to create 55 exemplar neurons representing the various morphological types that are reconstructed from the somatsensory cortex of a juvenile rat. The pipeline is then used to reconstruct a volumetric slice of a cortical circuit model that contains ∼210,000 neurons. The applicability of our pipeline to create highly realistic volumetric models of neocortical circuits is demonstrated with an *in silico* imaging experiment that simulates tissue visualization with brightfield microscopy. The results were evaluated with a group of domain experts to address their demands and also to extend the workflow based on their feedback.

**Conclusion** A systematic workflow is presented to create large scale synthetic tissue models of the neocortical circuitry. This workflow is fundamental to enlarge the scale of *in silico* neuroscientific optical experiments from several tens of cubic micrometers to a few cubic millimeters.

## Background

During the end of the last century, the neuroscience community has witnessed the birth of a revolutionary paradigm of scientific research: ‘*in silico* neuroscience’. This simulation-based approach has been established based on several aspects, fundamentally: the collection of sparse, yet comprehensive, experimental data to synthesize and build structural models of the brain in addition to the derivation of rigorous mathematical models that could interpret its function at different scales [KS98, GK09]. The integration between those structural and functional models is a principal key for reverse engineering and exploring the brain and gaining remarkable insights about its behavior [DVLMS06]. This approach has turned out to be a common practice first in domains where mathematical modeling is more evident, such as physics and engineering. In neuroscience, the term *in silico* appeared for the first time in the early 1990‘s when the community started to focus on computational modeling of the nervous system from the biophysical and circuit levels and up to the systems level [KS98]. Nevertheless, simulation-based research in neuroscience has not become widespread until more recently, when simulating complex biological systems has been afforded. This scientific revolution was a normal consequence of diversified factors including a huge quantum leap in computing technologies, a better understanding of the underlying principles of the brain and also the availability of experimental methods to collect the vast amounts of data that are necessary to fit the models [Mar06, ST07].

Understanding the complex functional and structural aspects of the mammalian brain relying solely on ‘wet’ lab experiments has been proven to be extremely limiting and time consuming. This is due to the fragmentation of the neuroscience knowledge; there are multiple brain regions, different types of animals models, distinct research scopes, and various approaches for addressing the same questions [MML*11]. The search space for unknown data is so broad, that it is debatable whether traditional experiments can provide enough data to answer all the questions in a reasonable time, unless a more systematic way is followed.

Integrating the *in silico* approach into the research loop complements the traditional *in vivo* and *in vitro* methods. Thanks to unifying brain models, *in silico* experiments allow the neuroscientists to efficiently test hypothesis, validate models and build in-depth knowledge as an outcome of the analysis of the resulting data from computer simulations [LO13, MMMF14, RCA*15]. Furthermore, these studies can also help to identify which pieces of unknown experimental data will provide the most information. The capacity of making new questions from *in silico* experiments establishes a strong link between theory and experimentation that would be very hard to do otherwise.

This systematic method can conveniently accelerate neuroscientific research pace and infer important predictions even for some experiments that are infeasible in the wet lab; for example due to the limited capability of the technology to probe a sample and measure variables or the physical impossibility of a manipulation such as silencing a specific cell type on a tissue sample or specimen. It also reduces the striking costs and efforts of the experimental procedures that are performed in the wet lab.

The reliability of the outcomes of an *in silico* experiment is subject to the presence of precise multi-scale models of brain tissue that could fit the conditions and the requirements of the experiment. In particular, the models that are relevant to this work are those which are biologically accurate at the level of organizational and electrophysiological properties of cells and their membranes.

Markram *et al*. presented a first-draft digital model of a piece—or slice— of the somatosensory cortex of a two-weeks old rat [RCA*15, MMR*15]. This model unifies a large amount of data from wet lab experiments and can reproduce a series of *in vitro* and *in vivo* results reported in the literature without any parameter tuning. However, the model is merely limited to simulating electrophysiological experiments. The fundamental objective of our work is focused on integrating further structural *volumetric* data into this model and extending its capabilities for performing *in silico* optical studies that can simulate light interaction with brain tissue.

We present a systematic approach for building realistic large scale volumetric models of the neocortical circuity from the morphological representations of the neurons; in which the model can account for light interaction with the different structures of the tissue. The models are created in three steps: meshing, voxelization, and data annotation (or tagging). To demonstrate the importance of the presented work, the resulting volumetric models are employed to simulate an optical experiment of imaging a cortical tissue sample with the brightfield microscope. This will allow us ultimately to establish comparisons between model and experimental results from different imaging techniques.

## Challenges and related studies

Structural modeling of neocortical circuits can be approached based on morphological, polygonal or volumetric models of the individual neurons composing the circuits. Each modeling approach has specific set of applications accompanied with certain level of complexity and limitations. Morphological models can be used to validate the skeletal representation of the neurons [PMA15], their connectivity patterns [KTvH*09] and their organization in the circuit [KSH*08], but they cannot be used, for example, for detailed visualization of electrophysiological simulations. Visualizing such spatiotemporal data requires highly detailed models that can provide multi-resolution, continuous and plausible representations of the neurons, such as polygonal mesh models [LHH*12, HSP12]. These polygonal models can accurately represent the cell membrane of the neurons, but they cannot characterize the light propagation in the tissue; they do not account for the intrinsic optical properties of the brain. Therefore, such models cannot be used to simulate optical experiments on a circuit level, for instance, microscopic [ABE*15a] or optogenetic experiments [NJGS13].

Simulating those experiments is constrained to the presence of detailed and multi-scale volumetric models of the brain that are capable of addressing light interaction with the tissue including absorption and scattering. There are also other *in silico* experiments, such as voltage sensitive dye imaging [CC10] and calcium imaging [SGHK03], that require more complicated models to simulate fluorescence. These volumetric models must be annotated with the actual spectral characteristics of the fluorescent structures embedded in the tissue to reflect an accurate response upon excitation at specific input wavelength.

In principle, volumetric models of the neurons can be obtained in a single step from their morphological skeletons using line voxelization [COK97]. However, the accuracy of the resulting volumes, in particular at the cell body and the branching points of the neurons, will be extremely limited. Moreover, addressing the scalability to precisely voxelize large scale neuronal circuits (micro-circuits, slice circuits or even meso-circuit) is not a trivial problem.

A correct approach of solving this problem entails creating tessellated polygonal meshes from the neuronal morphologies followed by building the volumes from the generated meshes using solid voxelization [ZGJ13, DCB*04]. Although convenient, this approach is not applicable in many cases because solid voxelization algorithms are conditioned by default to two-manifold or watertight polygonal meshes [Lla07]. Due to the complex structure of the morphological skeletons of the neurons and their reconstruction artifacts, the creation of watertight meshes from those morphologies is not an easy task. Polygonal modeling of neurons has been investigated in several studies for simulation, visualization and analysis purposes, but unfortunately they were not mainly concerned with the watertightness of the created polygonal meshes. This can be demonstrated in the work presented by Wilson *et al*. in Genesis [WBUB88], Glaser *et al*. [GG90] *in* Neurolucida and Gleeson et al. in neuroConstruct [GSS07]. These software packages have been designed solely for creating limited-quality and low level-of-detail meshes that can only fulfill their objectives. For instance, those created by Neurolucida were simplified to discrete cylinders that are disconnected between the different branches of the dendritic arbors as a result of the variations in their radii. This issue was resolved in neuroConstruct relying on tapered tubes to account for the difference in the radii along the branches, however, the authors have used uniform spheres to join the different branches at their bifurcation points. These meshes were watertight by definition, but they do not provide a smooth surface that can accurately reflect the structure of a neuron. Creating smooth and continuous polygonal models of the neurons has been discussed in two studies by Lasserre *et al*. [LHH*12] and Brito *et al*. [BMB*13a], but their meshes cannot be guaranteed to be watertight when the neuronal morphologies are badly reconstructed. Therefore, a novel meshing method that can handle the watertightness issues is strictly needed.

Building volumetric models of cortical tissue has been addressed in recent studies for the purpose of simulating microscopic experiments. Abdellah *et al*. have presented two computational methods for modeling fluorescence imaging with low- [ABE*15b, ABE*15a] and highly-scattering tissue models [ABE*17]. The extent of their volumetric models was limited to tiny blocks of the cortical circuitry in the order of tens to hundreds of cubic micrometers. Their pipeline has been used to extract a mesh block from the cortical column model by clipping each mesh whose soma is located within the spatial extent of this block and then convert those clipped meshes to a volume with solid voxelization. Before the clipping operation, the watertightness of each mesh in the block is verified. If the test fails, the mesh is reported and ignored during the voxelization stage. Consequently, this approach could limit the accuracy of any *in silico* experiment that utilizes their volumetric models. The algorithms, workflows and implementations discussed in the following sections are introduced to overcome these limitations and reduce a gap that is still largely unfulfilled.

## Contributions

1. Presenting an efficient meshing algorithm for creating piecewise watertight polygonal models of neocortical neurons from their morphologies.
2. Design and implementation of a scalable and distributed pipeline for creating polygonal mesh models of all the neurons in a given neocortical micro-circuit based on Blender [Ble16].
3. Design and implementation of a high performance solid voxelization software capable of building high resolution volumetric models of the cortical circuitry of few cubic millimeters extent.
4. Demonstrating the results with physically-based visualization of the volumetric models to simulate brightfield microscopic experiments.
5. Evaluating the results in collaboration with a group of domain experts and neuroscientists.

## Methods

Our approach for building scalable volumetric models of neuronal circuits from the experimentally reconstructed morphological skeletons is illustrated by Figure 1 and summarized in the following points:

1. Preprocessing the individual neuronal morphologies that compose the circuit to repair any artifacts that would impact the meshing process.
2. Creating smooth and watertight mesh models of the neurons from their morphologies.
3. Building local volumetric models of the neurons from their mesh models.
4. Integrating all the local volumes of the individual neurons into a single global volume dataset.
5. Annotating, or ‘tagging’ the global volumetric model of the circuit according to the criteria specified by the *in silico* study. For example, in clarified fluorescence experiments [TYHD14], the neurons will be tagged with the spectral characteristics of the different fluorescent dyes that are injected intracellularly. In optogenetic experiments, the volume will be tagged with the intrinsic optical properties of the cortical tissue [ABL*14] to account for precise light attenuation and accurate neuronal stimulation [BSM*11].

### Repairing morphological artifacts

The neuronal morphologies are reconstructed from imaging stacks obtained from different microscopes. These morphologies can be digitized either with semiautomated [OBSB07] or fully automated [HHC*03] tracing methods [HPD*08, GG90]. The digitization data can be stored in multiple file formats such as SWC and the Neurolucida proprietary formats [DSA02, TNTP08]. For convenience, the digitized data are loaded, converted and stored as a tree data structure. The skeletal tree of a neuron is defined by the following components: a cell body (or soma), sample points, segments, sections, and branches. The soma, which is the root of the tree, is usually described by a point, a radius and a two-dimensional contour of its projection onto a plane or a three-dimensional one extracted from a series of parallel cross sections. Each sample represents a point in the morphology having a certain position and the radius of the corresponding cross section at this point. Two consecutive samples define a connected segment, whereas a section is identified by a series of non-bifurcating segments and a branch is defined by a linear concatenation of sections. Figure 1-A illustrates these concepts.

**Figure 1:**
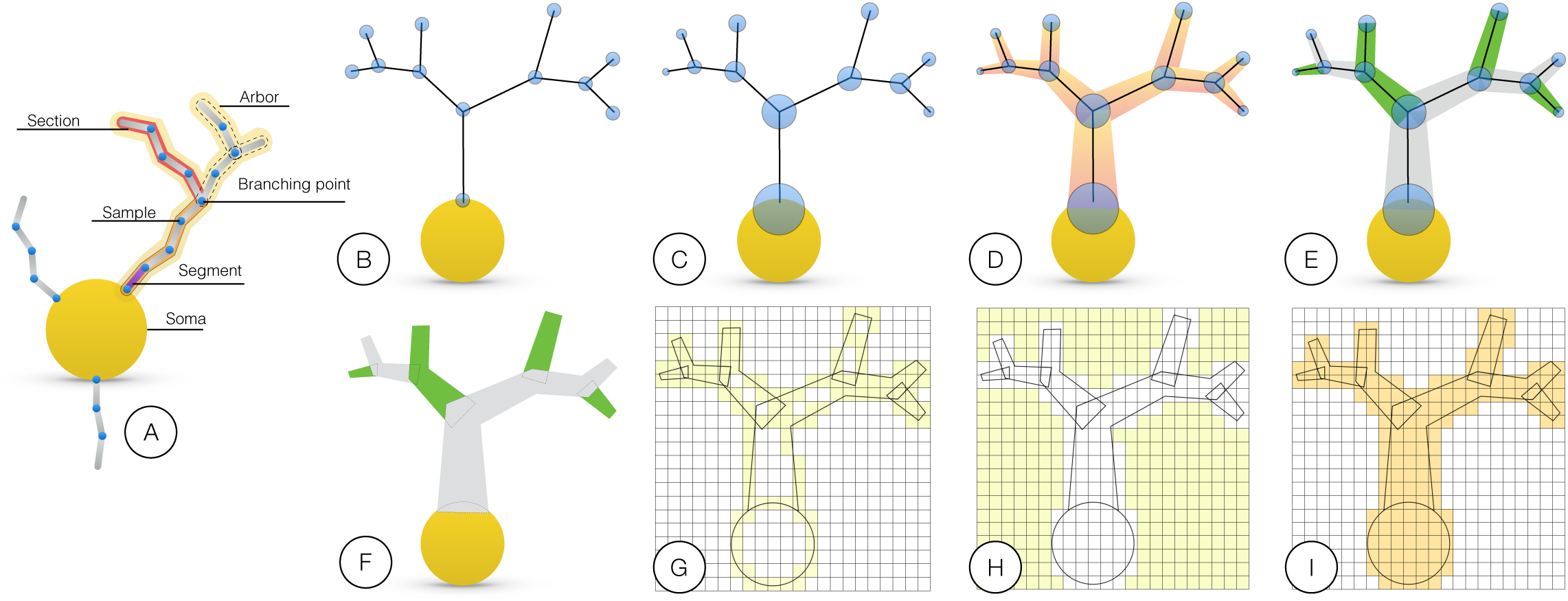
**An illustration of our proposed workflow for creating volumetric models of the neurons from their morphological skeletons**. (A) A graphical representation of a typical morphological skeleton of a neuron. To eliminate any visual distractions, the workflow will be illustrated using a single arbor sampled only at the branching points (B-F). The blue circles in (B) and (C) represent the positions of morphological samples of the neurons and the radii of their respective cross-sections. (D) The morphology structure is created by connecting the samples, segments, and branches together. (E) The primary branches that represent a continuation along the arbor (in the same color) are identified according to the radii of samples of the children branches at the bifurcation points. (F) The connected branches identified in (E) are converted into multiple mesh objects where each object is smooth and watertight. (G) The mesh objects are converted to intersecting volumetric shells with surface voxelization in the same volume. (H) Solid voxelization. The volume created in (G) is floodfilled to cover the extra-cellular space of the neurons. (I) The final volumetric model of a neuron is created by inverting the flood-filled volume to reflect a smooth, continuous and plausible representation of the neuron.

Due to certain reconstruction errors, morphologies can have acute artifacts that limit their usability for meshing. In this step, each morphological skeleton is investigated and repaired if it contains any of the following artifacts:

1. Disconnected branches from the soma (relatively distant); where the first sample of a first-order section is located far away from the soma.
2. Overlapping between the connections of first-order sections at the soma.
3. Intersecting branches with the soma; where multiple samples of the branch are located inside the soma extent.

These issues can severely deform the reconstructed three-dimensional profile of the soma, affect the smoothness of first-order branches of the mesh and potentially distort the continuity of the volumetric model of the neuron. The disconnected branches were fixed by repositioning the far away samples closer to the soma. The new locations of these samples were set based on the most distant sample that is given by the two-dimensional profile of the soma. For example, if the first order sample is located at 20 micrometers from the center of the soma, while the farthest profile point is located at 10 micrometers, then the position of this sample is updated to be located within 10 micrometers from the center along the same direction of the original sample.

The algorithm for creating a mesh for the soma is based on a deformation of an initial mesh into a physically plausible shape. Two branches influencing the same vertices of the initial mesh give rise to severe artifacts. Therefore, if two first-order branches or more overlap, the branch with largest diameter is marked to be a *primary* branch, while the others are ignored for this process. Finally, the samples that belong to first-order branches and are contained within the soma extent are removed entirely from the skeleton.

### Meshing: from morphological samples to polygons

In general, creating an accurate volumetric representation of a surface object requires a polygonal mesh model with certain geometrical aspects; the mesh has to be watertight, i.e. non intersecting, two-manifold [ED06]. Unfortunately, creating a single smooth, continuous and watertight polygonal mesh representation of the cell surface from a morphological skeleton is more difficult than it seems. Reconstructing a mesh model to approximate the soma surface is relatively simple, however, the main issues arise when (1) connecting first-order branches to the soma and (2) joining the branches to each others. Apart from the intrinsic difficulties, morphological reconstructions from wet lab experiments are not traced with membrane meshing in mind. Therefore, they may contain features and artifacts that can badly influence the branching process even if the artifacts are completely repaired. In certain cases, some branches can have extremely short sections with respect to their diameters or unexpected trifurcations that can distort the final mesh.

The existing approaches for building geometric representations of a neuron are not capable of creating a smooth, continuous and watertight surface of the cell membrane integrated into a single mesh object. In neuroConstruct, the neuron is modeled with discrete cylinders, each of them represents a single morphological segment [GSS07]. By definition, the cylinders are watertight surfaces, however, this technique underestimates the actual geometric shape of the branches. It introduces gaps or discontinuities between the segments that are not colinear. In contrast, the method presented by Lasserre et al. can be used to create high fidelity and continuous polygonal meshes of the neurons, but the resulting objects from the meshing process are not guaranteed to be watertight. Their algorithm resamples the entire morphological skeleton uniformly, and thus, the resampling step cannot handle bifurcations that are closer than the radii of the branching sections. Moreover, the somata are not reconstructed on a physically-plausible basis to reflect their actual shapes. This issue has been resolved by the method discussed by Brito et al. [BMB*13b]. They can also build watertight meshes for the branches, but their approach can be valid only if the morphological skeleton is artifact-free. The watertightness of the resulting meshes is not guaranteed if the length of the sections are relatively smaller than their radii or when two first-order branches are overlapping.

We present a novel approach to address the previous limitations and build highly realistic and smooth polygonal mesh models that are watertight ‘piecewise’. The resulting mesh consists of multiple ‘separate’ and ‘overlapping’ objects, where each individual object is continuous and watertight. In terms of voxelization, this piecewise watertight mesh is perfectly equivalent to a single connected watertight mesh that is almost impossible to reach in reality. The overlapping between the different objects guarantees the continuity of the volumetric model of the neuron, Figures 1-G and 1-I. The final result of the voxelization will be correct as long as the union of all the pieces provides a faithful representation of each component of the neuron. The mesh is split into three components: (1) a single object for the soma, (2) multiple objects for the neurites (or the arbors) and (3) (optionally) multiple objects for the spines if that information is available.

**Soma meshing** In advanced morphological reconstructions, the soma is precisely described by a three-dimensional profile that is obtained at multiple depths of field [PHT02]. In this case, the soma mesh object can be accurately created relying on the Possion surface reconstruction algorithm that converts sufficiently-dense point clouds to triangular meshes [KH13]. However, the majority of the existing morphologies represent the soma by a centroid, mean radius and in some cases a two-dimensional profile, and thus building a realistic soma object is relatively challenging [HPD*08].

Lasserre *et al*. presented a kernel-based approach for recovering the shape of the soma from a spherical polygonal kernel with 36 faces [LHH*12]. The firstorder branches of the neurons are connected to their closest free kernel face, and then the kernel is scaled up until the faces reach their respective branches. The resulting somata are considered a better approximation than a sphere, but they cannot reflect their actual shapes. Brito *et al*. have discussed a more plausible approach for reconstructing the shape of the soma based on mass spring system and Hook‘s law [BMB*13b, TPBF87, NMK*06]. Their method simulates the growth of the soma by pulling forces that emanate the first-order sections. However, their implementation has not been open sourced to reuse it.

We present a similar algorithm for reconstructing a realistic three-dimensional contour of the soma implemented with the physics library from Blender [Ble16, Ken15]. The algorithm simulates the growth of the soma by deforming the surface of a soft body sphere that is based on a mass spring model. The soma is initially modeled by an isotropic simplicial polyhedron that approximates a sphere, called icosphere [Ble11]. The icosphere is advantageous over a UV-mapped sphere because (1) the vertices are evenly distributed and (2) the geodesic polyhedron structure distributes the internal forces throughout the entire structure. As a trade-off between compute time and quality, the subdivision level of the icosphere is set to four. The radius of the icosphere is computed with respect to the minimal distance between the soma centroid and the initial points of all the first-order branches.

Each vertex of the initial icosphere is a control point and each edge represents a spring. For each first-order section, the initial cross-section is spherically projected to the icosphere and the vertices within this projection are selected to create a *hook modifier*, which is an ensemble of control points than remains rigid during the simulation. Before the hook is created, all the faces from the selected vertices are merged to create a single face that is reshaped into a circle with the same radius as the projected radius of the cross-section. During the simulation, each hook is moved towards its corresponding target section causing a pulling force. At the same time, the connecting polygons are progressively scaled to illustrated in Figure 2. If two or more first-order sections or their projections overlap, only the section with the largest diameter is considered. The other will be extended later to the soma centroid during the neurite generation.

**Figure 2:**
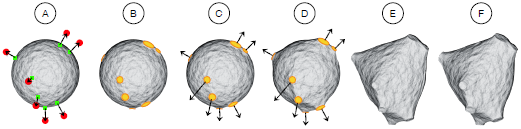
**Soma progressive reconstruction**. The soma is modeled by a soft body sphere in (A). The initial and final locations of the primary branches are illustrated by the green and red points respectively. The first-order sections are projected to the sphere to find out the vertices where the hooks will be created. The faces from each hook are merged into a single face and shaped into a circle (B). The hooks are pulled and the circles are scaled to match the size of the sections (C-E). The final soma is reconstructed in (F).

**Neurite meshing** To mesh a neurite, we first divide the morphology in a set of branches (concatenated non bifurcating sections) that span the entire morphological tree, Figure 1-E. The algorithm starts the first branch from the first-order section of the neurite. At the first bifurcation the section with the largest cross-section at the starting sample is chosen as the continuing section for the on-going branch, the rest are placed in a stack. The algorithm proceeds to the next bifurcation and repeats until a terminal section is reached. Once the branch is completed, the first section in the stack is popped and a new branch is created from there. The algorithm finishes when all sections have been processed.

Each branch is meshed separately using a poly-line and a circle bevel which is adjusted to the branch radius at each control point, Figure 1-F. The initial branch of each neurite is connected to the centroid of the soma with a conic section. For most branches this connection will not be visible, but it is necessary for those ones that were overlapping a thicker branch and did not participate in the soma generation. The whole algorithm requires only local information at each step so it runs very quickly and in linear time in relation to the number of sections.

### Voxelization: from polygonal to volumetric models

A straightforward approach to voxelize an entire neuronal circuit of a few hundred or thousand neurons is to create a polygonal mesh for each neuron in the circuit, merge all of them in a single mesh and feed that mesh into an existent robust solid voxelizer. However, this approach is infeasible due to the memory requirements needed to create the single aggregate mesh model of all neurons. We propose a novel and efficient CPU-based method for creating those volumetric models without the necessity of building joint models of neurons. We use a CPU implementation to not restrict the maximum volume data size to the memory of an acceleration device, *e.g*. a GPU [FC00, ED08, SS10]. To reduce the memory requirements of our algorithm, we use binary voxelization to store the volume (1 bit per voxel).

The volume is created in four steps: (1) computing the dimensions of the volume, (2) parallel surface voxelization for the piecewise meshes of all the neurons in the circuit, (3) parallel and slice-by-slice-based solid voxelization of the entire volume, and finally (4) annotating the volume.

The spatial extent of the circuit is obtained by transforming the piecewise mesh of each neuron to global coordinates, computing its axis-aligned bounding box, and finally calculating the union bounding box of all the meshes. The size of the volume is defined according to the circuit extent and a desired resolution. The volumetric shell of each component of the mesh is obtained with surface voxelization, Figure 1-G. This process rasterizes all the pieces conforming a mesh to find their intersecting faces with the volume. This step is easily parallelizable, as each cell can be processed independently. We only need to ensure that the set operations in the volume dataset are thread-safe.

Afterwards, the extracellular space is tagged by flood-filling the volume resulting from surface voxelization [BK93]. To parallelize this process, we have used a two-dimensional flood-filling algorithm that can be applied for each slice in the volume, Figure 1-H. and the final volume is created by inverting the flood-filled one to discard the intersecting voxels in the volume, Figure 1-I.

## Results and discussion

### Implementation

The meshing algorithm is implemented in the latest version of Blender (2.78) [Ble16]. The pipeline is designed to distribute the generation of all the meshes specified in a given circuit in parallel relying on a high performance computing cluster with 36 computing nodes, each shipped with 16 processors. The meshing application is configured to control the maximum branching orders of the axons and dendrites, control the quality of the meshes at various tessellation levels and to integrate the spines to the arbors if needed. This pipeline has been employed to create highly-tessellated and piecewise watertight meshes of the neurons that were defined in a recent digital slice based on the reconstructed circuit by Markram *et al*. [MMR*15]. This circuit (521× 2081× 2864 *μ*^3^) is composed of ∼210,000 neurons and spatially organized as seven neocortical column stacked together. Using 200 cores, all the meshes were created in eight hours approximately. On average, a single neuronal morphology is meshed in the order of hundreds of milliseconds to a few seconds. The meshes were stored according to the Stanford polygon file format (.ply) to reduce the overhead of reading them later during the voxelization process.

The voxelization algorithms (surface and solid) are implemented in C++, and parallelized using the standard OpenMP interface [DM98]. The quality of the resulting volumetric models is verified by inspecting the two-dimensional projections of the created volumes and comparing the results to an orthographic surface rendering image of the same neurons created by Blender.

### Physically-based reconstruction of the somata

To validate the generalization of the soma reconstruction algorithm, the meshing pipeline is applied to 55 exemplar neurons having different morphological types as described in [RCA*15, MMR*15]. The exemplars were carefully selected to reflect the diversity of the shapes of neocortical neurons. Figure 3 shows the eventual shapes of the reconstructed somata of only 20 neurons. The progressive reconstruction of all the 55 exemplars is provided as a supplementary movie [Abd17].

**Figure 3:**
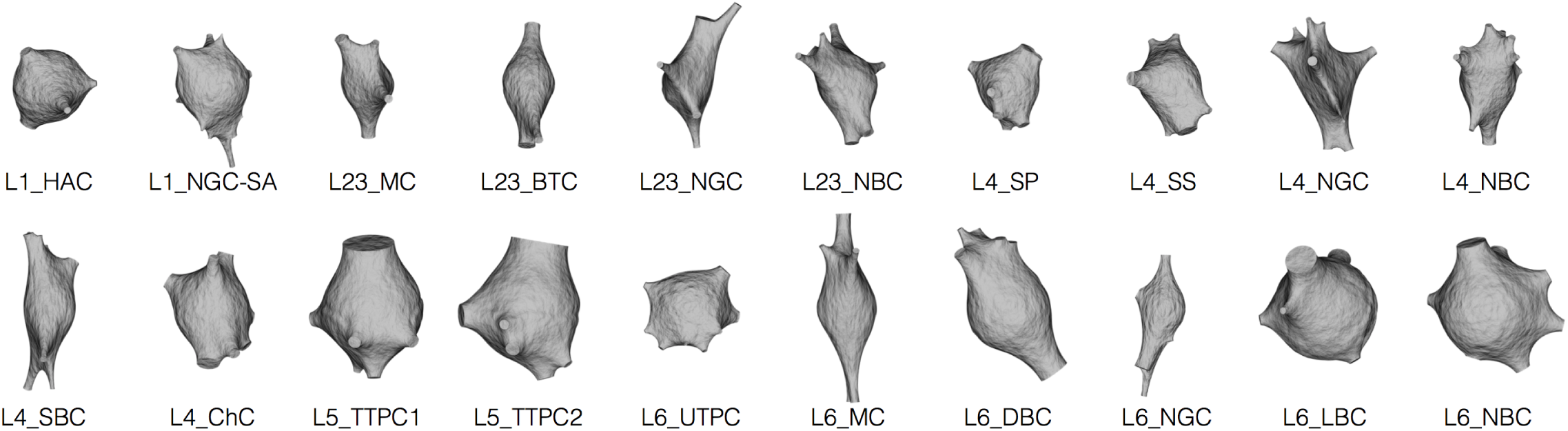
**Physically-plausible reconstruction of the somata of diverse neocortical neurons labeled by their morphological type**. The initial shape of the soma is defined by a soft body sphere that is deformed by pulling the corresponding vertices of each primary branch. The algorithm uses the soft body toolbox and the hook modifier in Blender [Ble16].

### Piecewise watertight polygonal modeling of the neurons

Figure 4 shows an exemplar piecewise watertight polygonal mesh of a pyramidal neuron generated from its morphological skeleton. Figure 4-C shows closeups of the meshes created for a group of other neurons having different morphological types. The resulting meshes of all the 55 exemplars are provided in high resolution in the supplementary files. The different objects of each mesh are rendered in different colors to highlight their integrity without being a single mesh object. The watertightness of the created meshes of the exemplar neurons was validated in MeshLab [CCR08]. All the neurons have been reported to have zero non-manifold edges and vertices.

**Figure 4:**
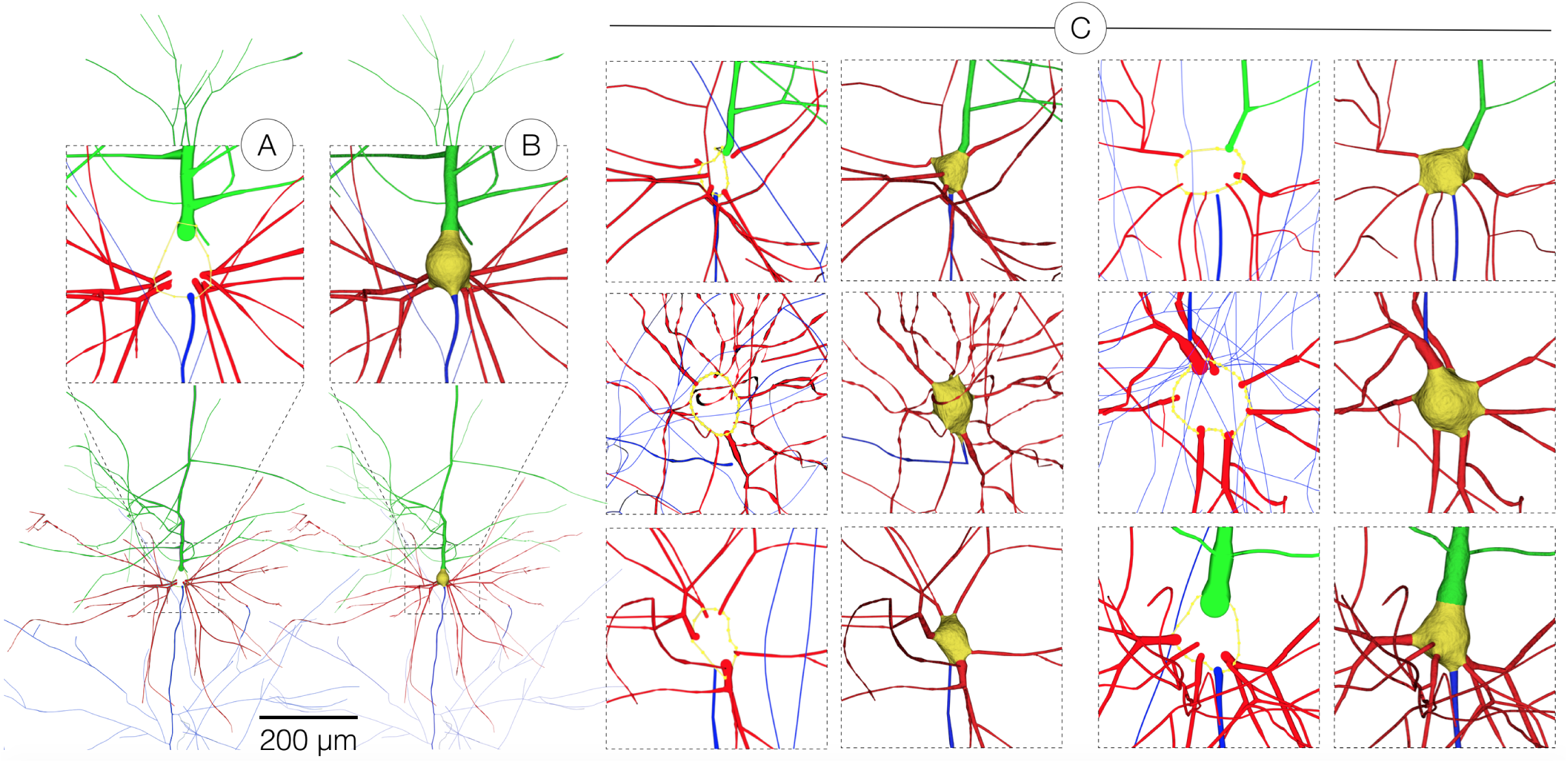
**Reconstruction of a piecewise watertight polygonal mesh model of a pyramidal neuron in (B) from its morphological skeleton in (A)**. In (C), the applicability of the proposed meshing algorithm is demonstrated with multiple neurons having diverse morphological types to validate its generality. The reconstruction results of the 55 exemplar neurons are provided in high resolution with the supplementary materials. The somata, basal dendrites, apical dendrites and axons are colored in yellow, red, green and blue respectively.

### Volumetric modeling of a neocortical circuit

The scalability of our voxelization workflow affords the creation of high resolution volumetric models of multi-level neocortical circuits (microcircuits, mesocircuits, slices) that are composed from a single neuron and up to an entire slice that contains ∼210,000 neurons. The target volume is created upon request from the neuroscientist according to his desired *in silico* experiment. Figure 5 illustrates the results of the main steps for creating an 8k volumetric model of a single spiny neuron from its mesh model. The volumetric shell of each component of the neuron is created with surface voxelization. The filling of the intracellular space of the neuron is done with solid voxelization to create a continuous and smooth volumetric representation of the neuron.

**Figure 5:**
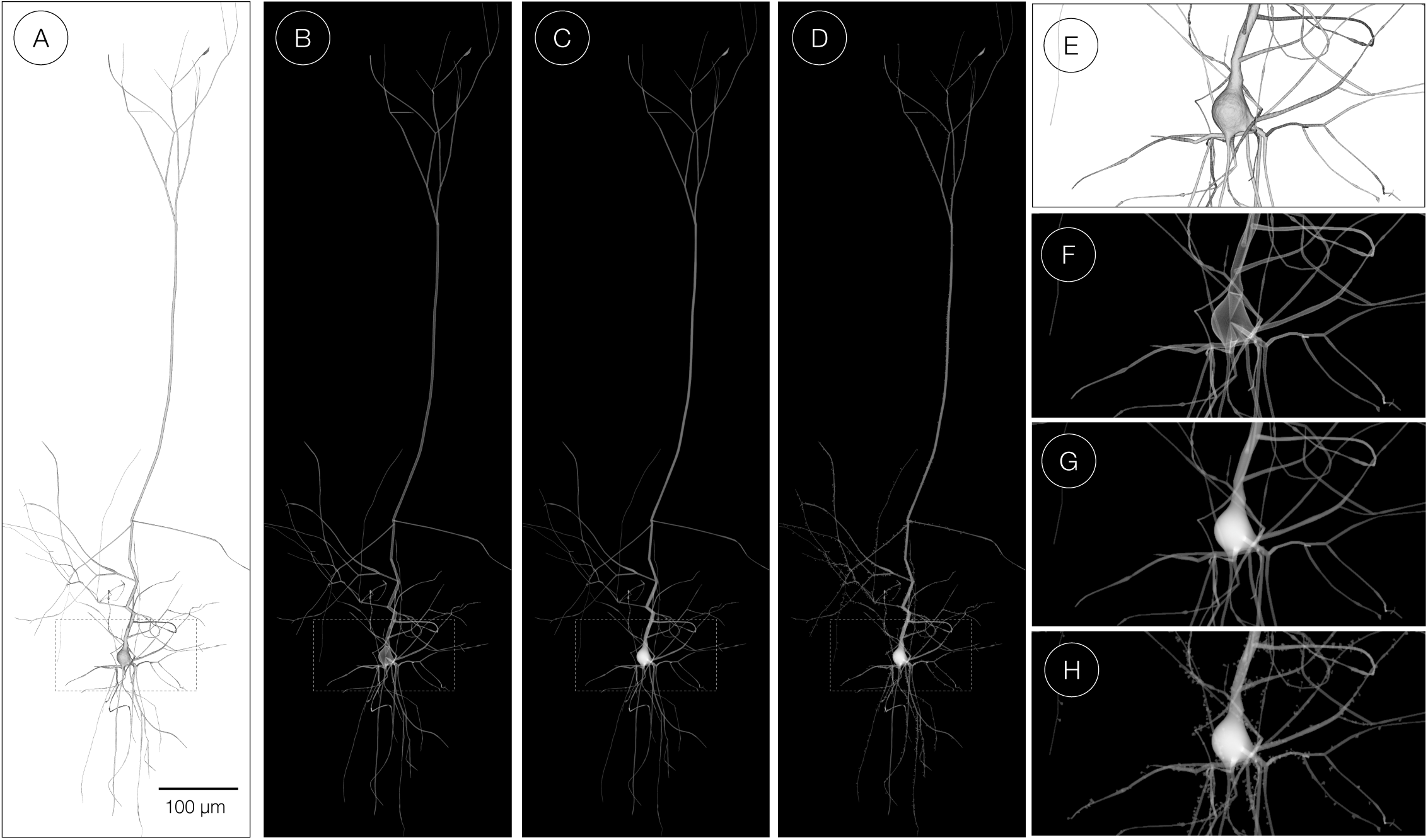
**The process of building a volumetric model of a single pyramidal neuron from its polygonal mesh**. The polygonal mesh model in (A) is converted to a volumetric shell with surface voxelization in (B) and a filled volume with solid voxelization in (C). In (D), the spines are integrated to the volume. The images in (E), (F), (G) and (H) are close ups for the renderings in (A), (B), (C) and (D) respectively. Notice the overlapping shells of the different branches and the soma that result due to the surface voxelization step in (F). In (G), the volume created with solid voxelization reflects a continuous, smooth and high fidelity representation of the entire neuron.

Figure 6 shows the results of volumizing multiple neocortical circuits with various scales that range from a single neuron and up to a slice circuit. Note that we only voxelize a fraction of neurons to be able to visualize the volume, but in principle the volumes were created for all the neurons composing the circuit. Referring to previous studies [ABE*15b, ABE*17], the scalability concerns addressed in this work has allowed the computational neuroscientists to extend the scale of their simulations from the size of the box colored in orange in Figure 6 to an entire slice.

**Figure 6:**
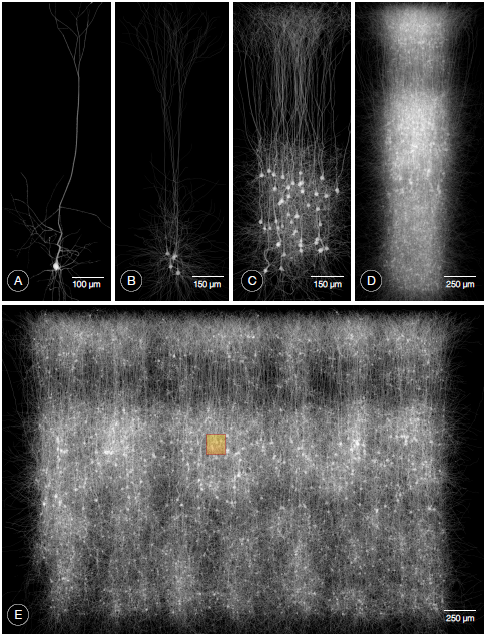
**Volumetric reconstructions of multiple neocortical circuits with solid voxelization**. The presented workflow is capable of creating large scale volumetric models for circuits with different complexity. (A) Single cell volume. (B) A group of five pyramidal neurons. (C) 5% of the pyramidal neurons that exist in layer five in the neocortical column. (D) 5% of all the neurons in a single column (containing ∼31,000 neurons). (E) A uniformlysampled selection of only 1% of the neurons in a digital slice composed of seven columns (containing ∼210,000 neurons) stacked together. The resolution of the largest dimension of each volume is set to 8000 voxels. The area covered by the orange box in (E) represents the maximum volumetric extent that could be simulated in similar previous studies [ABE*15b, ABE*17].

### Physically-plausible simulation of brightfield microscopy

To highlight the significance of this work, we briefly present a use case that utilizes the volumetric models created with our pipeline; a physically-plausible simulation of imaging neuronal tissue samples with brightfield microscopy. In general, this visualization is used to simulate the process of injecting the tissue with a specific dye or stain with certain optical characteristics to address the response of the tissue to this dye. Existing applications can use the models as well for performing other *in silico* optical studies such as [ABE*15b, ABE*17]. In this use case, the neurons are injected with Golig-based staining solution *in vitro*. Then, the sample is scanned with inverted brightfield microscope at multiple focal distances to visualize the neuronal connectivity and the in-focus structures of the neurons. We developed a computational model of the brightfield microscope that can simulate its optical setup and would allow us to perform this experiment *in silico*. For this purpose, a circuit consisting of only five neurons is volumized and annotated with the optical properties of the Golgi stain. Moreover, the virtual light source used in the simulation uses the spectral response of a Xenon lamp. The results of this *in silico* experiment is shown in Figure 7. The microscopic simulation is implemented on top of the physically-based rendering toolkit [PH10, PH12].

**Figure 7:**
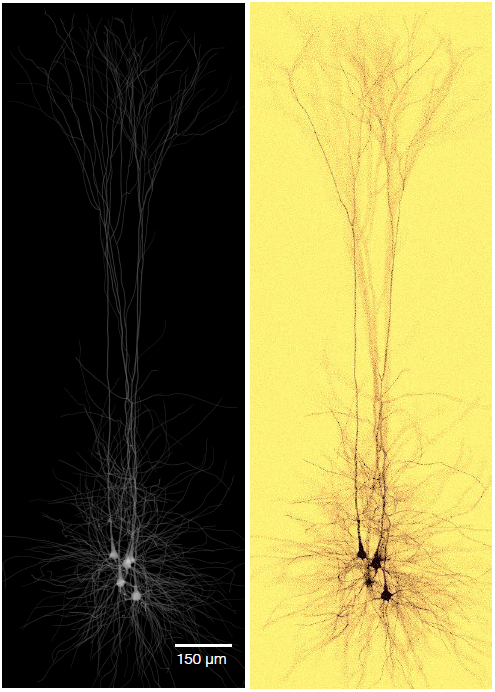
**In silico imaging of neuronal tissue with brightfield microscope**. The volumetric model (left) is annotated with the optical characteristics of Golgi‘s silver stain. The *in silico* image created in (B) is used to study the visual response of different dyes used in the *in vitro* experiment.

### Workflow evaluation

The significance of the results was discussed in collaboration with a group of domain experts including neurobiologists and computational neuroscientists. We requested their feedback mainly on the following aspects: the plausibility of the volumes of the 55 exemplar neurons, their opinions about the simulation of the brightfield microscope and the scalability of the workflow. They agreed that the neuronal models of the different exemplars, in particular the somata, are much more realistic than the current models they use in their experiments. They were also impressed with the rendering in Figure 7 saying that it is really hard to discriminate from those they have seen in the wet lab. They also suggested to use this optical simulation tool to experiment and validate the result of using other kinds of stains with different optical properties. They were also extremely motivated to see the results of other *in silico* experiments that simulate fluorescence microscopes and in particular for imaging brainbows [CCL*13] where each neuron is annotated with different fluorescent dye. Concerning the scalability, they expressed their deep interest to integrate our workflow into their pipeline to be capable of creating larger circuits. We have also received several requests to extend the pipeline for building volumetric models of other brain regions, for example the hippocampus, and also for reconstructing different types of structures such as neuroglial cells and vasculature.

## Conclusions and perspectives

We presented a novel and systematic approach for building large scale volumetric models of the neocortical circuitry of a two-week old Juvenile rat. An efficient and configurable pipeline is designed to convert the neuronal morphologies into smooth and high fidelity volumes without the necessity to create connected watertight polygonal mesh models of the neurons. The morphologies are repaired in a preprocessing step and then converted into piecewise watertight polygonal mesh models to build realistic volumetric models of the brain tissue with solid voxelization. The pipeline has been employed to create high resolution volumes for multiple neocortical circuits with a single neuron and up to a slice circuit that contains ∼210,000 neurons. The entire pipeline is parallelized to afford the voxelization of huge circuits in few hours, which was totally infeasible in the past. The results were discussed collaboratively with a group of experts to evaluate their plausibility. The significance of the presented method is demonstrated with a direct application for simulating the imaging of cortical tissue with brightfield microscopy.

We are currently working on improving the performance of the voxelization workflow to allow the creation of the volumes on distributed computing nodes. We are also considering building a web-based interface for the entire pipeline that can facilitate its usability in particular for pure biologists who have limited programing experiences. We will investigate the capability of extending the workflow to generate volumes for hippocampal neurons, neuroglial cells and vasculature to address the requests of the neuroscientists.

## Software and data availability

The source code is available from the corresponding author on a reasonable request. The neocortical circuits and the morphologies are available on-line at https://github.com/BlueBrain.

## Supplementary materials

1. A high resolution movie showing the somata reconstructed with our blender-based mesh generation software for the 55 exemplars cells selected from the neocortical column [Abd17].
2. High quality renderings of the generated 55 exemplar meshes and their morphologies.

## Authors’ contributions

MA convinced the study, designed and implemented the meshing and voxelization workflows and drafted the manuscripts. JH contributed to the discussions of the meshing and the voxelization algorithms. NA contributed to the design of the meshing workflow in Blender. JH, NA and SE contributed to discussions and suggestions to complete the manuscript. HM and FS supervised the project. All the authors read and approved the final manuscript.

## Authors details

Blue Brain Project (BBP), École Polytechnique Fédérale de Lausanne (EPFL), Biotech Campus, Chemin des Mines 9, 1202 Geneva, Switzerland.

## Declarations

This publication was supported in part by the Blue Brain Project (BBP), the Swiss National Science Foundation under Grant 200020-129525 and the King Abdullah University of Science and Technology (KAUST) through the KAUST-EPFL alliance for

## Acronyms

**CPU** Central Processing Unit

**GPU** Graphics Processing Unit

